# Testing the priming effect in the deep ocean: are microbes too starved to consume recalcitrant organic carbon?

**DOI:** 10.1101/2025.06.13.658936

**Authors:** Richard LaBrie, Corday R. Selden, Nagissa Mahmoudi

## Abstract

Deep ocean dissolved organic carbon (DOC) is one of the largest pools of reduced carbon on Earth. Many DOC compounds escape microbial degradation and persist for thousands of years in the ocean. Although many hypotheses have been proposed, the mechanisms responsible for this long-term stability remain unresolved. Heterotrophic microorganisms in the deep ocean are energetically starved and exhibit low metabolic activity. Here, we investigated whether the severe energy limitation in deep sea environments acts as a barrier to microbial degradation of DOC. We hypothesized that alleviating this energetic barrier through the addition of labile compounds (i.e., the priming effect) could stimulate microbial consumption of DOC. We conducted 62-day bottle incubations with deep seawater from the Southern Ocean that were amended with simple organic carbon, nitrogen- and/or phosphorus-containing compounds. We tracked DOC concentration, cell abundance and microbial community structure over the course of the experiment. Our results show no evidence of a priming effect regardless of the priming compound. However, priming compounds selected for distinct microbial populations with little overlap among amended bottles even when compounds were chemically similar. *Pseudoalteromonas* and *Pseudomonas* were enriched across all amended bottles, and their competition for labile substrates likely contributed to observed variations in DOC consumption. Our results reveal that the persistence of DOC is not driven by the energetic state of deep-sea microbes. In contrast, our results suggest that inputs of fresh carbon to the deep ocean are more likely to increase DOC sequestration, via the microbial carbon pump, rather than stimulate DOC consumption.

**IMPORTANCE:** The oceans store vast amounts dissolved organic carbon (DOC) that can resist microbial degradation for thousands of years. The mechanisms that underlie the long-term stability of DOC in the ocean are still debated. Microorganisms in this environment exhibit low metabolic activity and are energetically starved. We tested whether the microbial degradation of DOC could be stimulated through the addition of labile compounds. Surprisingly, alleviating energetic constraints did not stimulate the consumption of deep ocean DOC. Additionally, our results suggest that competition among taxa is an important constraint on dissolved organic carbon consumption, with implications for ecosystem processing. Our study indicates that an increase in fresh organic carbon to the deep ocean may enhance carbon sequestration since marine microbes are known to produce recalcitrant compounds. Among other applications, this finding is of consequence for ongoing geoengineering efforts that aim to remove atmospheric carbon by increasing carbon export to the deep sea.

## INTRODUCTION

Marine dissolved organic carbon (DOC) represents one of the largest pools of reduced carbon on Earth, comparable in size to the carbon contained in the atmosphere (1). In the deep ocean, most of the DOC pool is considered to be recalcitrant, meaning that it escapes microbial degradation and can persist for millennia (2). Despite decades of research, the precise mechanisms that underlie the long-term stability of DOC in the deep ocean remain unresolved. One explanation attributes its low biological reactivity (i.e., bioavailability) to the intrinsic chemical properties of the compounds that make up the pool (3–5). For example, major classes of compounds found within the DOC pool include carboxyl-rich alicyclic molecules in which fused alicyclic rings contribute to their low bioavailability (6), and carotenoid degradation products (7)—how these molecules resist microbial degradation is unknown. Alternatively, the ‘dilution hypothesis’ proposes that individual compounds within the DOC pool occur at such low concentrations that they are effectively inaccessible to microbes (8–10). Supporting this view, mass spectrometry analyses have revealed that deep ocean DOC contains over than 100,000 distinct compounds, most present at concentrations below 10 pmol L^−1^ (11). This suggests that if individual compounds are replenished, microbes could consume them until their concentrations again fall below detectable thresholds, thereby sustaining a dynamic equilibrium within the DOC pool.

While chemical composition and dilution may both contribute to the persistence of DOC in the ocean, we explore a third perspective centered on the energetic state of deep-sea microbes. Readily available (i.e. labile) carbon substrates are scarce in the deep ocean such that microbial communities are severely energy-limited (12, 13). Consequently, many microbes enter a state of metabolic dormancy or drastically reduced metabolic activity (14–17). For example, experimental studies using seawater from the North Atlantic Ocean observed limited enzymatic response to complex DOC by bathypelagic (1,000–4,000m deep) microbes—up to 2 orders of magnitude lower—compared to surface and mesopelagic (200–1,000m deep) depths (14). This indicates that microbes in deeper waters had substantially lower levels of metabolic activity potentially due to the limited supply of fresh carbon to these depths (14, 15, 17).

One potential mechanism for overcoming this energetic limitation is through the addition of labile carbon compounds which can stimulate microbes to degrade more complex, recalcitrant organic compounds—known as the priming effect. This phenomenon was first discovered in soil science a century ago (18), and it has only been recently considered in aquatic sciences (19). Priming studies are usually conducted using simple carbon compounds such as glucose (20, 21); however, organic compounds containing nitrogen (dissolved organic nitrogen i.e., DON) and/or phosphorus (dissolved organic phosphorus i.e., DOP) may be more effective as they not only provide carbon and energy but also essential nutrients that are in low concentrations in organic forms in the deep ocean (22–24).

In this study, we investigated whether microbial degradation of deep ocean DOC could be primed by the addition of labile compounds, particularly by the addition of DOP and DON. We hypothesized that these energetically rich and readily available compounds could alleviate the energetic and nutritional constraints of deep-sea microbes, thereby stimulating them to consume and degrade DOC. To test this, we conducted 62-day laboratory incubations using Southern Ocean seawater collected at 2500 m in which microbial communities were primed with various simple organic compounds. We tracked DOC concentrations and cell abundance to assess the consumption of the added priming compounds and background DOC. In addition, we assessed microbial community structure at the onset of incubations and after 28 days to determine if priming led to shifts in the composition and diversity of microbial communities. Our results suggest that the stability of DOC is not driven by the energetic state of deep-sea microbes and that growth dynamics between microbial taxa shape the consumption of DOC.

## MATERIALS AND METHODS

### Seawater collection

Seawater was collected aboard the *RRS Discovery* in January 2024 in the South Atlantic sector of the Southern Ocean (56.41601°S, 0.573025°W, **Fig. S1**). We identified the source seawater for these experiments as Circumpolar Deep Water. In addition to the station location and depth, this appraisal is based on the following water mass characteristics (25): the sampled water was cold and salty, with an *in situ* temperature and salinity of −0.12°C and 34.662, respectively. Moreover, dissolved oxygen concentrations were relatively low (225.0 μmol kg^−1^) and nutrient concentrations were relatively high (nitrate = 33.5 ±0.2 μM, phosphate = 2.30 ±0.01 μM; silicate = 128.1 ±0.02 μM), consistent with an aged water mass. Circumpolar Deep Water is formed at depth in the Southern Ocean where deep waters from the North Atlantic, Indian and Pacific Oceans combine (25, 26). Due to this origin, Circumpolar Deep Water is characteristically old, with typical radiocarbon ages indicating ∼1,400 years since ventilation (27).

Seawater was collected at a depth of 2500 m using Niskin bottles mounted to a steel rosette equipped with a conductivity, temperature, depth (CTD, Seabird Electronics 911plus), dissolved oxygen sensor (SBE-43). Seawater was transferred to dark acid-washed (10% HCl, 1 day) high-density polyethylene carboys using acid-washed (10% HCl, 1 day) food-grade tubing. Carboys were stored in the dark at 4°C for the remainder of the voyage (∼4 weeks), then packaged with ice packs designed for extended storage (ICEpack Xtreme) and shipped from Walvis Bay, Namibia, to McGill University, Montréal, Canada, in a heavy-duty cool box. Upon arrival at McGill University, carboys were immediately stored at 4°C in the dark until processing the following day.

### Bottle Incubations

Prior to incubation, seawater was sequentially filtered through 3 µm and 0.8 µm polycarbonate filter (Millipore, USA) using a peristaltic pump system to remove large particles (**Fig. 1**). This allowed us to specifically target DOC and bacterioplankton during our experiments. Approximately 1 L volume of filtered seawater was transferred to 1.2 L combusted glass bottles. The priming effect was tested using four distinct amendments: (1) glucose was added to represent a labile carbon substrate; (2) amino acids were added as proxy for DON; (3) glucose-6-phosphate was added as a proxy for DOP; and (4) a combination of both amino acids and glucose-6-phosphate were added together to assess the cumulative impact of DON and DOP. Priming solutions were prepared by dissolving glucose, or amino acids (L-arginine, L-aspartic acid, L-glutamic acid and L-serine, each comprising ∼25 mol%) or glucose-6-phosphate in 100 mL of milli-Q water. A total volume of 10 mL of priming solution was added to increase total DOC concentration by 0.9 mg/L.

**Figure 1.**
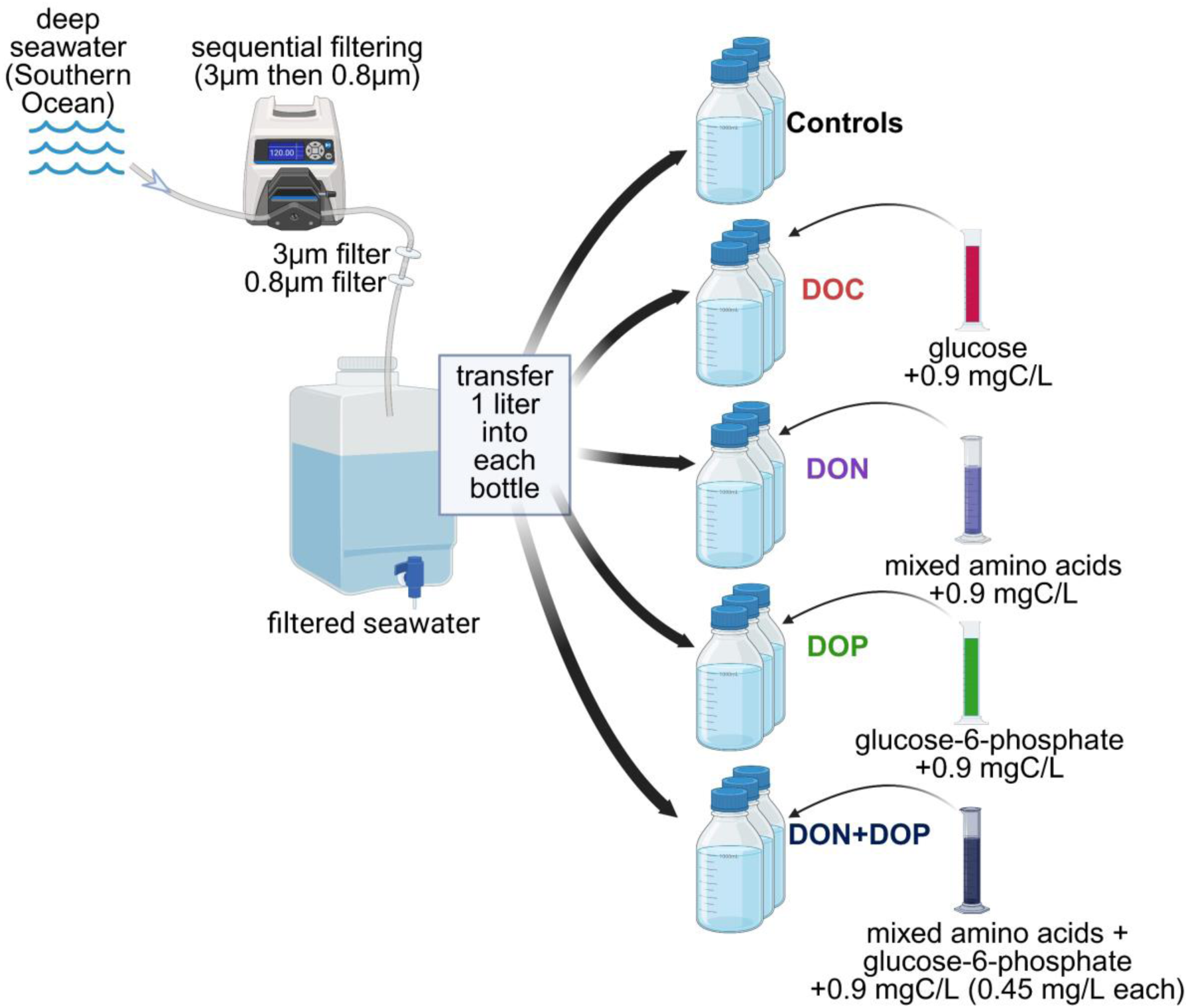
Schematic of the experimental setup used in this study to test the priming effect in the deep ocean.

This concentration ensured a measurable difference between amended bottles and no-amendment control experiments. In the case of the DON+DOP-amended bottles, an equal amount of the amino acids and glucose-6-phosphate solutions was added to the filtered seawater so that the total DOC concentration increased by 0.9 mg/L. All controls and amendments were done in triplicate such that there was a total of 15 bottles, including three control bottles with filtered seawater, three with glucose, three with DON, three with DOP and three with both DON and DOP. All bottles were kept at 4°C in the dark for a total of 62 days.

Sampling was conducted daily for 5 days (day 0 to day 4), followed by additional sampling on days 7, 10, 14, 21, 28 and 62. Subsamples were collected by transferring seawater from each bottle into a combusted, pre-conditioned glass beaker to avoid contamination. Subsamples were analyzed for DOC, DON and DOP concentrations, cell counts, and microbial community composition (only on the initial timepoint and day 28 due to water limitations).

### DOC, DON and DOP quantification

Subsamples were filtered on 0.2 µm polycarbonate filters using a vacuum hand-pump (Cole-Parmer). DOC was measured on a high temperature catalytic oxidation (HTCO) TOC-Vcpn analyzer (Shimadzu) equipped with a TNM-1 module for nitrogen measurement at Institut des Sciences de la Mer (ISMER), Canada. Briefly, samples were acidified to pH < 2 using concentrated HCl (12 M, Fisher Scientific ACSPlus) and stored in pre-combusted (450 °C for 5 h) glass vials at 4°C. Total dissolved nitrogen (TDN) was measured using the same subsample. DOC and TDN concentrations were validated using deep seawater reference (28).

DON was calculated as the difference between dissolved inorganic nitrogen (DIN, NO_x_ [nitrite + nitrate], ammonium) and TDN. Samples for nutrients (DIN and phosphate) were filtered 0.2µm polycarbonate filters and stored frozen (−20°C) until analysis on a SAN++ continuous flow (Skalar, Neatherlands) using colorimetric [NO_x_] or fluorometric ammonium] methods at CERC OCEAN, Dalhousie University, Canada. Detection limits were 0.01 µmol L^−1^ for nitrite, 0.05 µmol L^−1^ for nitrate, and 0.06 µmol L^−1^ for ammonium.

DOP was calculated as the difference between phosphate and total dissolved phosphorus (TDP). Samples for TDP were filtered and stored at −20°C until they were sent out for analysis at the University of Hawai’i Hilo Analytical Lab. TDP was analyzed using a Lachat Quikchem 8500 Flow Injection Analysis System (Hach, USA) with a detection limit of 0.25µmol L^−1^. Samples for phosphate were measured concurrently with inorganic nitrogen species, and the detection limit was 0.01 µmol L^−1^.

### Cell counts

Samples for microbial cell abundance were fixed with 0.1% final concentration glutaraldehyde grade I, flash-frozen, and kept at −80°C until analysis. Samples were stained with SYBR green 1 (1% final concentration) in tris-EDTA buffer (10 mM tris and 1 mM EDTA) (29) for 10 min in the dark and counted on a CytoFLEX flow cytometer (Beckman Coulter, USA) at ISMER.

### Microbial Community Analysis

Microbial community composition was characterized using the 16S rRNA amplicon sequencing at day 0 and 28. A total of 400mL of water was filtered at each time point using 0.2 µm sterile polycarbonate filters. Filters were kept at −80°C until further processing. DNA was extracted from filters using a SOX kit (Metagenom Bio Life Science, Canada) following manufacturers’ protocol and quantified using a Nanodrop One spectrophotometer (Thermo Scientific, USA). DNA amplification, library preparation and Illumina sequencing of 16S rRNA genes were done at the Integrated Microbiome Resources at Dalhousie University (30). Briefly, a dual-indexing, one-step PCR was done using a Nextera XT v2 kit (Illumina) with the adapters and index provided by the kit. Region V4-V5 of the 16S gene was targeted with forward primer 515FB (GTGYCAGCMGCCGCGGTAA) and reverse primer 926R (CCGYCAATTYMTTTRAGTTT) (31). Sequencing reads were analyzed with the DADA2 pipeline following the tutorial v.1.16 (32) using the R software v4.3.1 (33). After processing, 489,535 reads were retained (83%) representing 189 amplicon sequence variants (ASV). These were then taxonomically classified based on the SILVA database v138. One control sample was removed from the analysis due to low read count, and “mitochondria” was removed from the dataset; other samples were rarefied to 15,000 reads.

To assess differences in microbial community structure and diversity across amendments, we used non-metric multidimensional scaling (NMDS) to characterize beta-diversity using the *vegan* package (34) and calculated richness using the *vegan* package and Faith’s phylogenetic diversity (Faith’s PD, 35) using the *picante* package (36) to assess alpha-diversity. Statistical significance was tested using permutational analysis of variance (PERMANOVA, adonis2 function in *vegan*), pairwise Wilcoxon tests, and differences in microbial community structure among bottles were assessed using similarity profile (SIMPROF) analysis (37). All data analyses were performed on R Studio (33).

## RESULTS AND DISCUSSION

### No evidence of the priming effect found on deep ocean DOC

Our experiments were designed to test whether the introduction of labile compounds stimulated the breakdown of deep ocean DOC that is typically considered to be recalcitrant. Since we added carbon-based priming compounds, our starting concentration of DOC was higher in our amended bottles compared to our control bottles. If priming occurred, we expected the final DOC concentrations in amended bottles to be lower than those in control bottles, reflecting consumption of deep ocean DOC over the course of the experiment. In contrast to our hypothesis, we found no evidence of a priming effect on deep ocean DOC across all amended bottles (**Fig. 2A**). Instead, DOC concentrations in all amended bottles reached the same concentration as the control bottles by day 62 (DOC_final_ = DOC_control_, ANOVA, p-value > 0.05), indicating that the labile compounds that were added were consumed by microbial communities (DOC_final_ < DOC_initial_, ANOVA, p-value < 0.05), but that consumption of the background DOC pool was negligible.

**Figure 2.**
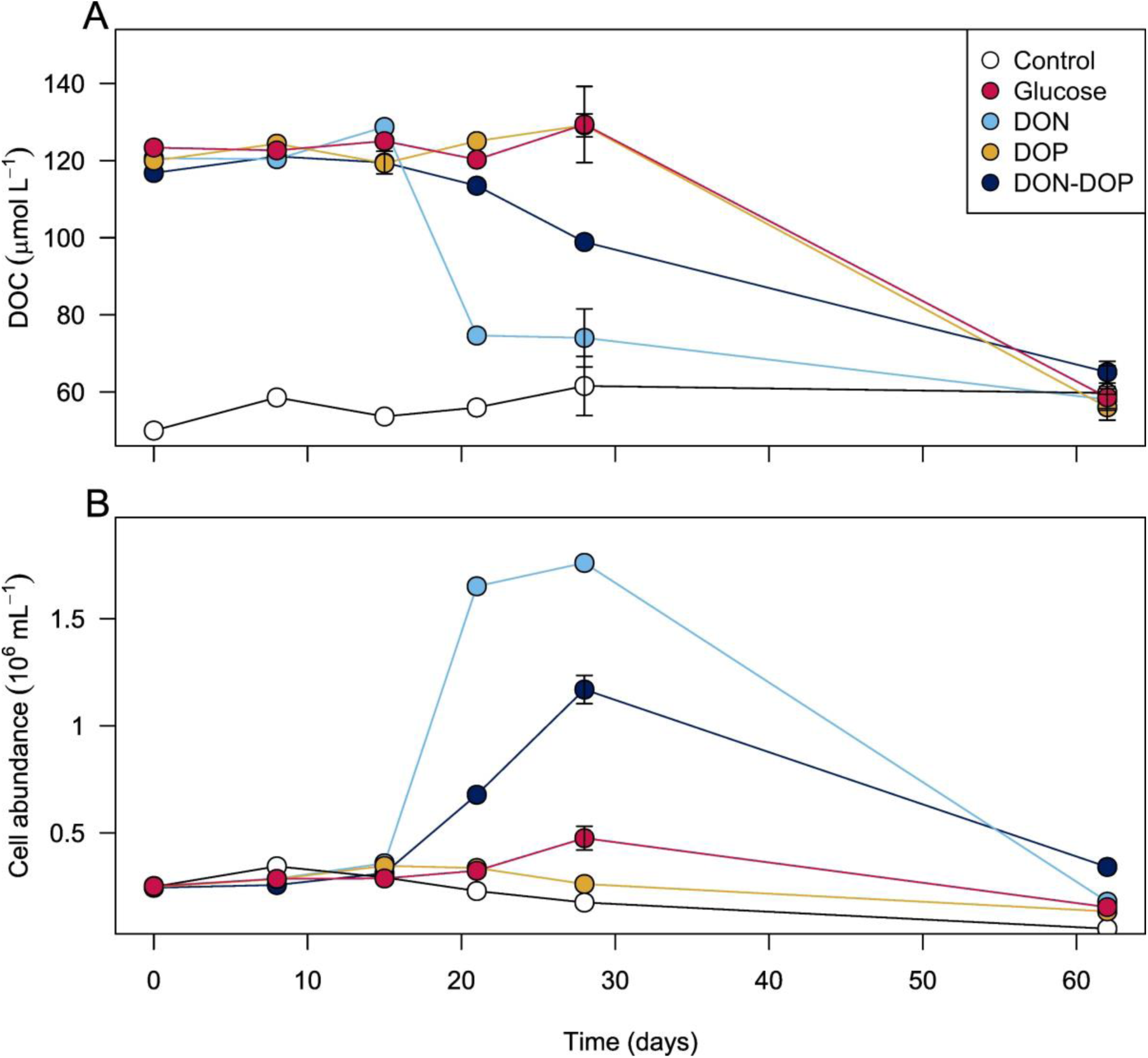
Temporal variations in DOC concentration (A) and cell abundance (B) over the course of bottle incubations using deep seawater from the Southern Ocean. Error bars represent standard error (SE) among treatment triplicates. For some sampling points, the error was smaller than the data point.

While the priming effect is a well-documented phenomenon in soil science (38), its occurrence in aquatic ecosystems is more controversial, with mixed results in both sediments and the water column. Priming was observed in North Sea sediments, where the addition of labile phytoplankton DOC enhanced consumption of background DOC by up to 31% (39). Conversely, no evidence of priming was found in sediments from the North-East Atlantic using similar methodology (40). In contrast, experiments in the Sargasso Sea showed significant surface DOC uptake after the addition glucose and inorganic nutrients (20). As there were no evidence of DOC degradation in unamended waters, Carlson et al., (20) concluded that semilabile DOC—the fraction that persists for 1.5 years (2)—was consumed in their amended bottles. In another study, filtered lake water was kept in the dark at 4°C for 12 years, ensuring that all remaining DOC was recalcitrant; subsequent addition of labile compounds did not stimulate microbial uptake of RDOC (21). In our study, the addition of labile compounds was not sufficient to stimulated the degradation deep ocean DOC, in which semilabile compounds are considered exhausted (2). Taken together, these results suggest that the reactivity of the DOC pools matters. While the input of labile compounds can prime microbial degradation of more reactive DOC, our results suggest that this effect has limits and does not extend to the most recalcitrant carbon pools on Earth.

### Differential consumption and utilization of labile substrates

Although all our amended bottles reached background DOC concentrations by day 62, there was preferential consumption among the added priming compounds (**Fig. 2A; Fig. S2**). Interestingly, we observed a long lag period in all our amended bottles, such that our added priming compounds persisted for ∼2 to 3 weeks prior to being consumed. Previous work used radiotracers to estimate the turnover time of glucose in the deep Southern Ocean at 80 to >365 days and concluded that most of the deep ocean microbial community was inactive or dormant, i.e., in an state of low metabolic activity (41). This suggests that deep sea microbial communities may require a period of acclimation before becoming active, as observed in other bathypelagic communities (14, 17).

The DON bottles showed the fastest consumption (38% decrease by day 21; ∼65% of added C), suggesting a higher affinity for amino acids compared to the other added compounds. This was followed by the DON+DOP mixture which showed a 15% decrease by day 28 (∼27% of added C). In the case of the glucose-amended bottles, there was no detectable decrease in DOC by day 28. We anticipated a rapid consumption of DOP in our amended bottles since DOP is cycled rapidly throughout oceanic depths (24, 42). As the deep ocean is replete in inorganic phosphorus compared to the surface ocean (>2µM), DOP is thought to be used as a source of energy and carbon for deep sea microbes (24, 43). Moreover, the deep ocean is highly limited in DOP compared to DOC with ratios of ∼3500C:1P (23), suggesting a high affinity of deep sea microbes for phosphorus-containing compounds (44, 45). Yet, we observed that DOP was not consumed for over 28 days while most of the DON (i.e., amino acids) were consumed by day 21. This implies that the microbial communities in our incubations could use amino acids more efficiently, which could indicate greater starvation of nitrogen than phosphorus. Nevertheless, the DOP was eventually fully consumed as DOC reached background concentration by day 62.

In all amended bottles, cell abundance dynamics closely followed DOC consumption patterns (**Fig. 2B**). The most striking increase in cell abundance was in the DON-amended bottles with a seven-fold increase after 28 days, reaching a maximum of ∼1.8×10^6^ cells mL^−1^. Such high cell abundance is typical of blooms in the surface ocean (46, 47) during which DOC is not a limiting factor for microbial growth. This indicates that microbial communities could efficiently use amino acids to build new cell biomass. Once the added compounds were exhausted, the microbial population returned to in situ abundance, similar to dynamics observed in the surface ocean (48). In the DON+DOP bottles, there was a five-fold increase in cell abundance after 28 days of incubation. It is possible that we missed the sampling window for peak cell abundance for these bottles since DOC concentrations were still high by day-28. In the glucose-amended bottles, cell abundance nearly doubled by day 28 implying that some glucose was consumed, but not enough to measure a decrease in DOC concentration. Given that DOC concentrations reached background levels in the glucose and DOP-amended bottles by day 62 indicating that these added priming compounds were in fact consumed, we suspect there was an increase in cell abundance that corresponded to this consumption. However, our sampling time points likely missed the exact time during the incubation period when cell abundance increased such that abundances remained relatively stable at 3.0 ± 1.0 x10^5^ and 2.7 ± 0.8×10^5^ cells mL^−1^, respectively (**Fig. 2B**). Both synchronous (49) and asynchronous (50) responses between cell abundance and DOC consumption have been observed in experimental studies with deep sea water.

### Addition of labile compounds caused reproducible shifts in microbial community structure

Although the added compounds were not fully consumed in the first month, we observed a reproducible impact on microbial diversity and community structure by day 28. The NMDS ordination plot revealed significant differences between amended and control bottles (PERMANOVA, p-value<0.01, **Fig. 3A**). The taxonomic composition of microbial communities clustered tightly according to the amendments and there was no overlap among them (**Fig. 3A**). This indicates that the addition of the priming compounds exerted a strong selection pressure on the observed community structure.

**Figure 3.**
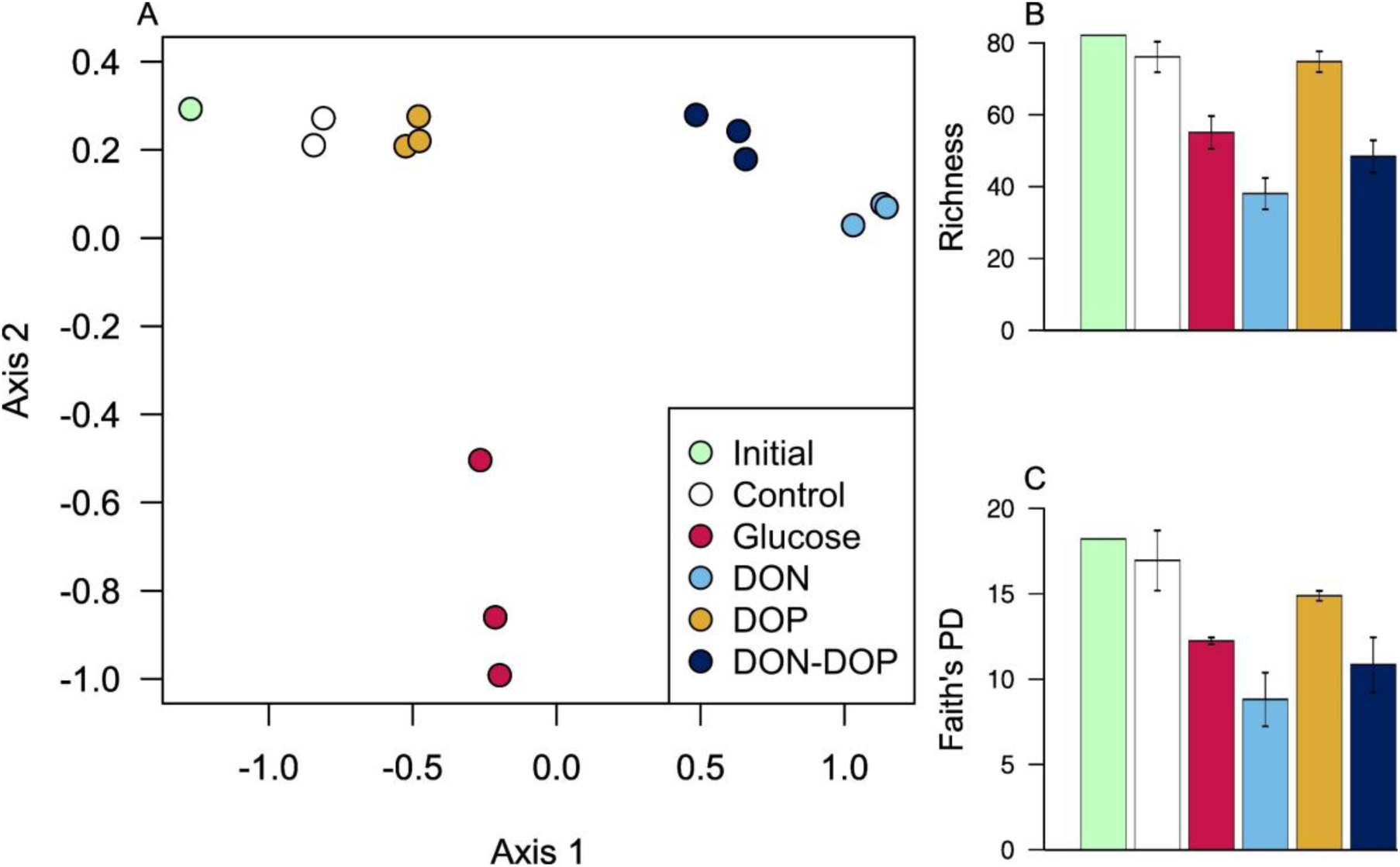
Analysis of microbial communities using NMDS ordination of Bray-Curtis dissimilarity matrices showing no overlap among amended bottles and controls (A). Priming compounds generally decreased the number of taxa in the community (B) and Faith’s phylogenetic diversity (C), with larger decrease in DON-, DON+DOP- and glucose-amended bottles. The NMDS and barplots share the same legend.

As expected, the taxonomic composition of the control bottles slightly differed from the initial seawater (SIMPROF, p-value<0.01, **Fig. 3A**, **Fig. 4A**). This is a phenomenon coined the bottle effect (e.g., 51, 52), where changes in community composition may happen in just a few days (53). In our case, this shift was comparatively smaller than in other studies (54, 55), suggesting that the observed changes in amended bottles were more related to the addition of the priming compounds rather than bottle confinement. Interestingly, the taxonomic composition of the DOP-amended bottles on day 28 remained similar to the starting community while the other amendments greatly differed (**Fig. 4A**, pairwise PERMANOVA, p-value<0.01). Likewise, the cell abundance in the DOP-amended bottles at day 28 was still low and similar to those observed in the control bottles (**Fig. 2B**), indicating that both the community composition and cell numbers had not substantially changed by this time.

**Figure 4.**
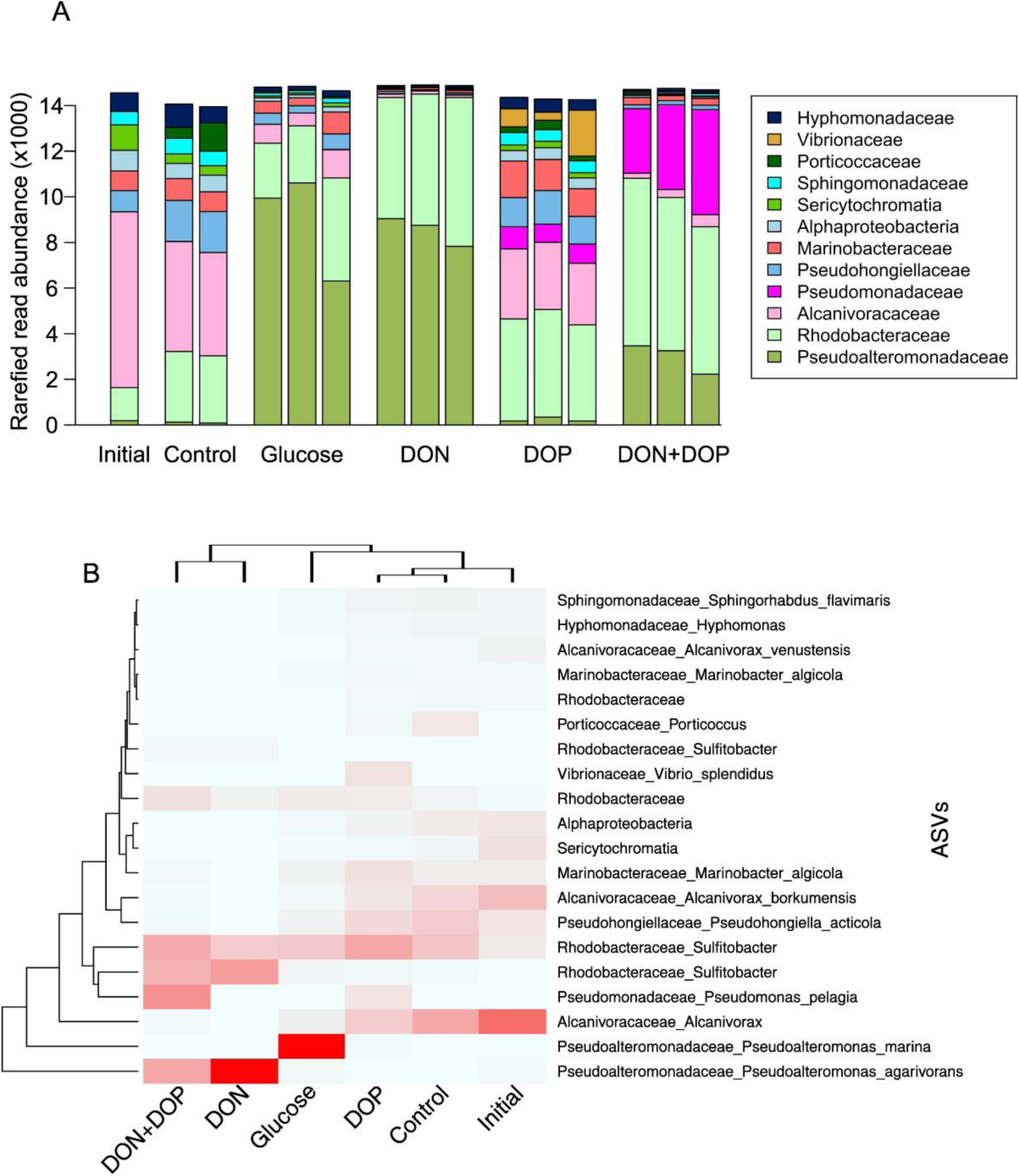
Taxa representing more than 1% of total read counts across control and amended bottles (A) and heatmap of the 20 most abundant ASVs across amendments where darker colors represent higher relative abundance across all treatments (B). The initial seawater community was sampled at the start of the experiment; other samples were collected after 28 days. Taxa are represented at the family level, except for two that are at the class level (Alphaproteobacteria and Sericytochromatia). Note that the Y-axis in panel A goes up to 14,000 and that samples were rarefied to 15,000 reads.

Microbial diversity decreased in the amended bottles which indicates that priming compounds enriched for a few populations that became dominant over time. Both richness (total number of observed AVSs) and Faith’s PD were lower in amended bottles compared to the controls (Wilcoxon test, p-value<0.05, **Fig. 3B**, **3C**). The largest decrease in diversity was observed in DON-amended bottles, followed by DON+DOP and glucose bottles, thus closely following cell abundance dynamics. These changes were not statistically significant using multiple comparison test (Wilcoxon test, p>0.05), but were generally significant when compared individually (t-test, p-value<0.05), suggesting a lack of statistical power rather than truly similar richness and diversity among amendments. The exception to this was the DOP-amended bottles which had similar diversity on day 28 as the control bottles (Wilcoxon test or t-test, p-value>0.34), suggesting that the selection pressure was lower compared to the other amended bottles.

Across all bottles, microbial communities were dominated by *Gammaproteobacteria* (60% of total read abundance) and *Alphaproteobacteria* (37%). In the initial seawater, microbial communities at the family level were dominated by *Alcanivoraceae*, *Rhodobacteraceae* and *Pseudohongiellaceae*, together representing 67% of total read abundance (**Fig. 4A**). These taxa were still abundant in control and DOP-amended bottles by day 28. However, in the other amended bottles (DON, DON+DOP and glucose) most of the initial taxa were outcompeted and community composition had drastically changed by day 28 (pairwise PERMANOVA, p-value<0.01, **Fig. 4A**). Moreover, most of the abundant taxa in DON-, DON+DOP- and glucose-amended bottles were either rare taxa representing less than 1% of read abundance (56), or undetected in the initial community (**Fig. S3**). This indicates that the addition of DON and glucose enriched for taxa that were likely dormant or in low abundance and subsequently increased in numbers once their preferred substrate became available. In general, each priming compound appeared to promote the growth of specific microbial populations, consistent with previous experimental studies (52, 53). In some cases, there was little to no overlap in these groups even though the added priming compounds were chemically similar. For example, *Pseudoalteromonas marina* was dominant (∼60%) in the glucose bottles (**Fig. 4B**) but represented only a minor proportion (< 1%) of the community in the DOP-only bottles that were amended with glucose-6-phosphate.

Interestingly, when both DON and DOP were added we observed an intermediate lag in DOC consumption—longer than in DON-only bottles but shorter than in those amended with DOP alone. We suspect that the presence of both priming compounds (DON and DOP) enriched two distinct microbial populations that were subsequently forced to compete for resources. In the DON-amended bottles, *Pseudoalteromonas agarivorans* was dominant (**Fig. 4B**), and rapidly consumed the added priming compounds (i.e. mixed amino acids). However, when both DON and DOP were present, there was an enrichment of other taxa including *Pseudomonas pelagia* which was also enriched in the DOP-only bottles. The coexistence and competition of these two populations may have slowed the growth of *P. agarivorans*, resulting in a lag in DOC consumption (**Fig. 2A**). In contrast, in the DOP-only bottles, the relative abundance of *P. pelagia* was lower which suggests that it grows faster through co-metabolism of DON and DOP than with DOP alone. Likewise, the lack of observed enrichment of *P. agarivorans* in the DOP only bottles indicates that it requires DON for growth. These results suggest microbial interactions appeared to strongly influence the timing and extent of DOC consumption across our amended bottles.

It was hypothesized that complex DOC compounds may favor cooperative behavior in microbial communities (57, 58) whereas simple compounds may instead increase competition among taxa as specialized enzymes are not required to acquire simple compounds. While complex compounds may be found in surface waters and in sediments, the deep ocean DOC pool consists of small molecules (59–61). As such, competition for substrate may be common in the deep ocean. Moreover, this pattern in microbial behaviour—cooperation versus competition—could also be a mechanism explaining the absence of priming in environments where DOC is less reactive. Taken together, these results suggest that carbon dynamics in deep sea environments may be intimately related to microbial community structure and interactions among microbes.

## CONCLUSIONS

The causes for the long-term stability of deep ocean DOC are still debated, with some arguing that compounds could be too dilute to be consumed by microbes (8, 10), and others that some compounds may simply be intrinsically recalcitrant to microbes due to their chemical structure (3, 62). Here, we tested whether the priming effect could remove the energetic barrier that deep microbes face, thus potentially stimulating the consumption of DOC. Our results do not support a priming effect, regardless of the priming compound added. These findings suggest that persistence of deep ocean DOC is not driven by the energetic or metabolic state of deep-sea microbial communities. In contrast, the opposite may be true: fresh DOC inputs to the deep ocean may induce more carbon sequestration through the microbial carbon pump as microbes are known to produce recalcitrant compounds from simple compounds (50, 63). Collectively, our work suggests that input of labile compounds in the deep ocean through sinking particles and viral lysis (64, 65) is more likely to result in increased DOC sequestration via the microbial carbon pump than net degradation through the priming effect. Likewise, ongoing geoengineering efforts which aim to increase carbon export to the deep ocean (66) will likely not stimulate deep ocean DOC degradation which could have dampened carbon sequestration efforts if priming were to occur.

## ACKNOWLEDGEMENTS

We would like to thank the captain and crew of the *RRS Discovery* and the science party of cruise DY172, without whom this work would not have been possible. In particular, we thank Dr. Tom Bibby (University of Southampton, UK) for supporting the collection of seawater for this study. We also thank Dr. Nina Schuback (Swiss Polar Institute, Chelsea Technologies Ltd.) for helping CRS lug 40 L of sample water across Walvis Bay, Namibia. This work was supported by a NSERC Discovery (RGPIN-2019-04228) grant and New Frontiers in Research Fund (NFRF) Exploration grant (NFRFE-2019-00794) to NM as well as McGill Wares and Trottier Space Institute postdoctoral fellowship to RL. The field work, including the water mass characterization, was supported by the Natural Environment Research Council (NERC) (grant NE/W000903/1) to Drs. Tom Bibby, Mark Moore, and Maeve Lohan (University of Southampton, UK).

## DATA AVAILABILITY

All data necessary to produce the results and figures in this paper are available on github.com/RichardLaBrie/SO_Priming. Raw sequences were deposited on NCBI Sequence Read Archive (accession number PRJNA1274019)

