## Supplementary figures for "Testing the priming effect in the deep ocean: are microbes too starved to consume recalcitrant organic carbon?"

### ADDITIONAL FIGURES

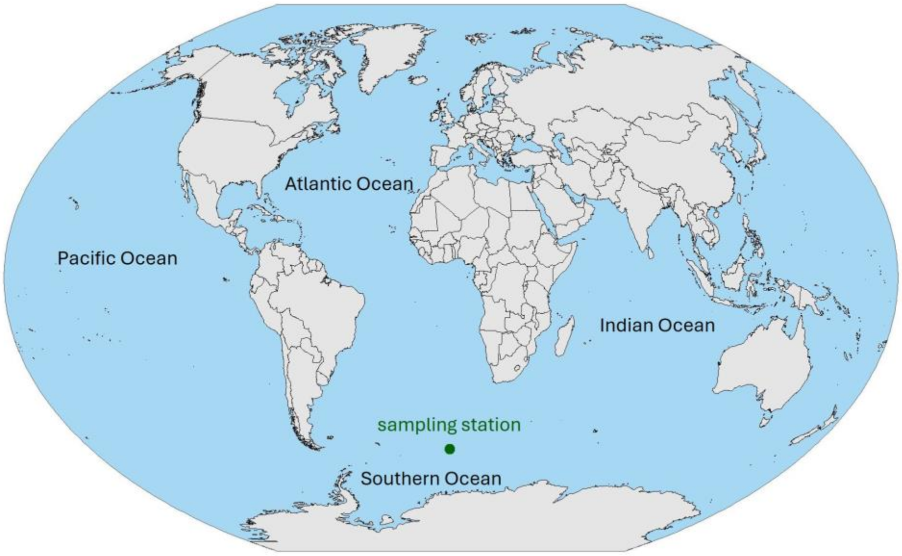


**Figure S1**. Sampling location for seawater used for bottle incubations.

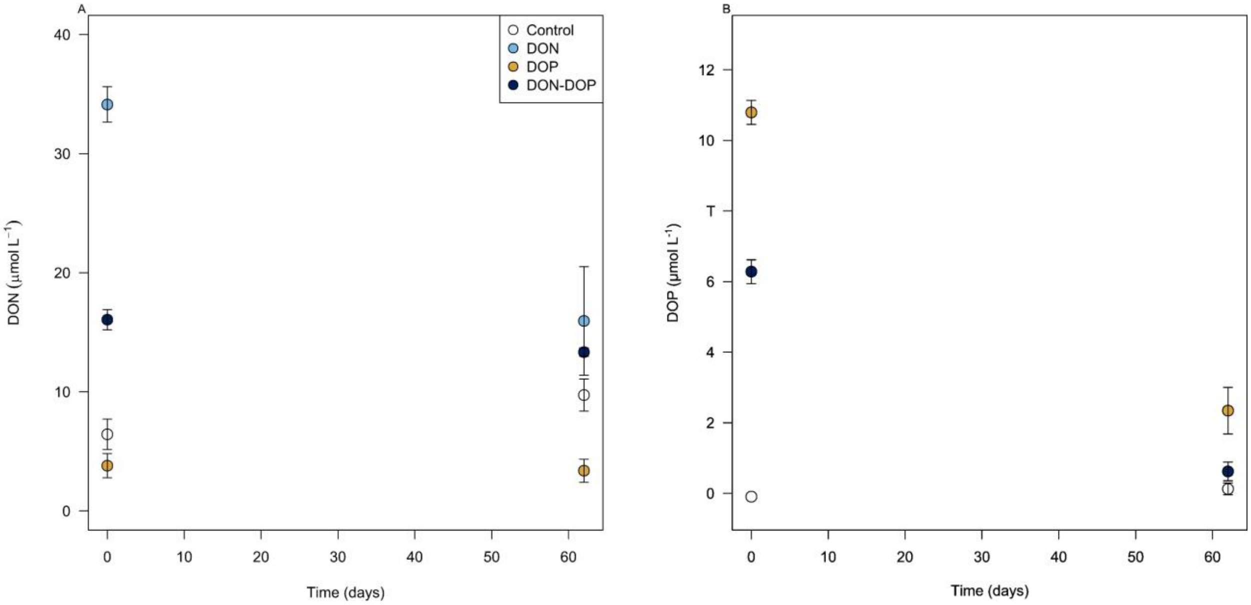


Figure S2. Concentrations of dissolved organic nitrogen (A) and dissolved organic phosphorus (B) in the initial and final time points. This confirmed that the decrease in DOC was attributed to consumption of the added compounds and not from background DOC. The difference in DON concentrations between amended bottles and controls is likely attributed to urea, which is a likely end product of amino acids degradation.

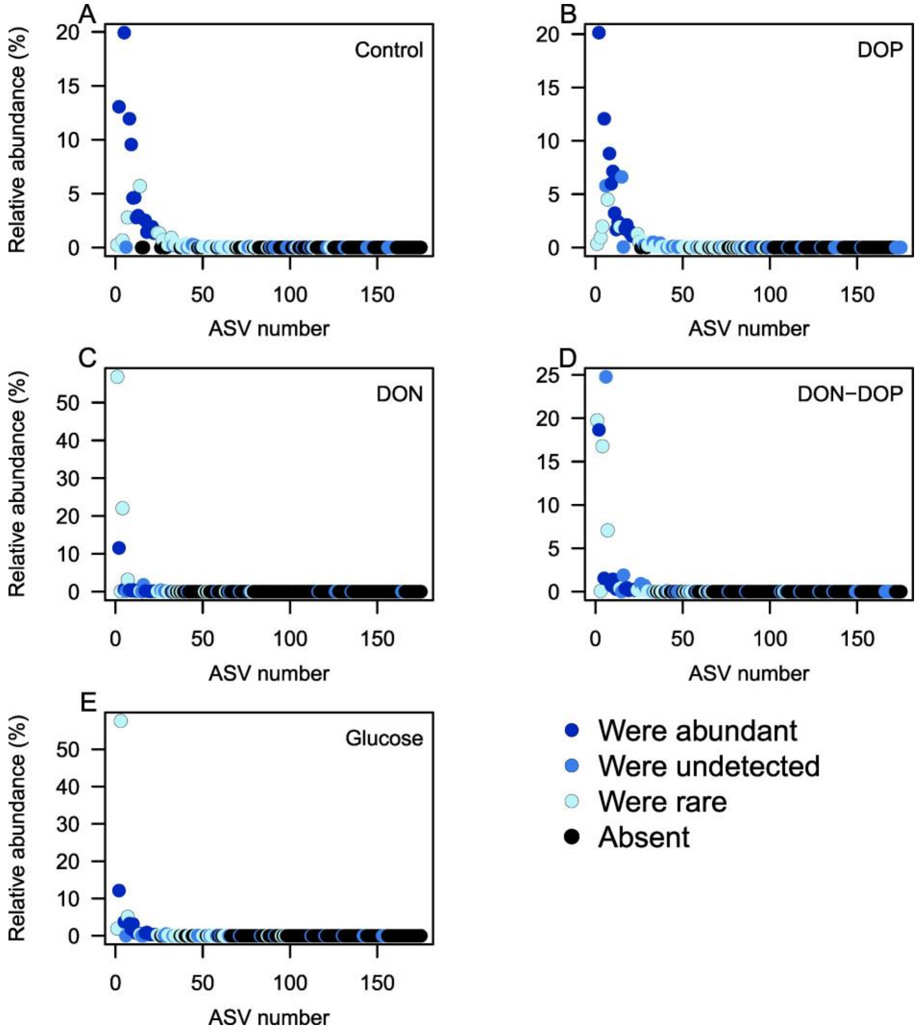


Figure S3. ASVs ranked as a function of their relative abundance across all bottles and represented as relative abundance in their respective communities (indicated on the right of the plots). Points are coloured as a function of their relative abundance in the initial time point, except for black. Dark blue: ASV was abundant (>1%) in the initial community; blue: ASV was not detected in the initial community; light blue: ASV was rare (<1%) in the initial community; black: ASV is not detected in the community on day 28.
